# Global distribution and host specificity of plastid-bearing microeukaryotes across coral holobionts

**DOI:** 10.64898/2026.10.20.759601

**Authors:** Joana Krause-Massaguer, Bradley A. Weiler, Javier del Campo

## Abstract

Corals form intricate symbiotic relationships with diverse microbial communities that influence their physiology, evolution, and ecological function. In 16S rRNA gene microbiome analyses, sequences taxonomically assigned to chloroplasts are often discarded as noise, yet they contain valuable ecological information. Here we present a meta-analysis of publicly available 16S rRNA gene datasets examining the diversity and global distribution of plastid-bearing microeukaryotic symbionts in corals. We reprocessed 2,679 samples from 45 studies, generating a dataset of 10,950 amplicon sequence variants (ASVs) from 13 million read pairs across 73 coral species, 23 families, all major oceanic regions, and three host health states. Corallicolida and *Ostreobium* were highly abundant across all coral lineages, health states, and locations, confirming them as the most widespread plastid-bearing coral symbionts after the Symbiodiniaceae. The most abundant ASVs exhibited strong host specificity, with most sequences restricted to specific coral lineages, a pattern consistent with phylosymbiosis, and supported across the full dataset by ASV-level community profiles. Phylogenetic placement of apicomplexan reads classified most of them as Corallicolida and revealed ASVs consistent with a novel apicomplexan-related lineage, tentatively designated ARL-XXI. Six distinct *Ostreobium* clades were recovered, three described here for the first time, with the North West Pacific emerging as a potential hotspot of *Ostreobium* diversity. Together, these results highlight the value of retaining plastid-derived sequences and underscore the importance of microeukaryotic-inclusive molecular surveys for understanding the ecological dynamics of the coral holobiont.

## Introduction

Corals are key species for ecosystem structure and function, supporting highly diverse benthic communities by providing refugia, breeding grounds, and food sources for a wide range of marine species. These ecosystem functions are also mediated by diverse and abundant microbial communities (archaea, bacteria, fungi, protists, and viruses) that, together with the coral host, form the coral holobiont [1]. Microbial members of the holobiont play roles in photosynthesis, biogeochemical cycling, nutrient supply, and pathogen suppression, and are suggested to influence the evolution, physiology, and ecological functions of the host [2, 3]. Despite the critical role of microorganisms, most research on coral symbioses has historically focused on bacteria. Microbial eukaryotes (protists and unicellular fungi) constitute a substantial fraction of coral microbial communities yet receive considerably less attention than prokaryotes, a bias reflected in genomic databases (80% bacterial vs. 20% eukaryotic sequences) [4]. The best characterized coral protist symbionts are dinoflagellates from the family Symbiodiniaceae, particularly in reef-building corals. Beyond the Symbiodiniaceae, however, a broad range of protistan symbionts remains largely unexplored, including their diversity and ecological roles [5, 6]. These include phototrophic microeukaryotes such as chlorophytes and rhodophytes, as well as heterotrophic microeukaryotes, including apicomplexans, fungi, ciliates, labyrinthulomycetes, syndiniales, and other dinoflagellates.

Two of the most widespread and abundant non-Symbiodiniaceae protists in corals are Corallicolida and *Ostreobium*. Corallicolida have been found in ∼70% of corals worldwide [7]. They are members of the Apicomplexa, a large subphylum of parasitic alveolates that include causative agents of human disease. Corallicolida, like their dinoflagellate counterparts, live intracellularly within the gastrodermis of coral hosts [8]. They have an apicoplast that, although non-photosynthetic, retains genes involved in chlorophyll synthesis. Although found in healthy individuals and showing no evidence of pathogenicity, Corallicolida’s evolutionary placement among obligate parasites raises questions about their functional role within the holobiont [7]. Corallicolida have been found in high abundance in *Paramuricea clavata* colonies highly susceptible to heat stress, suggesting that they play a role in coral fitness and thermal resilience [9].

*Ostreobium* is a green, filamentous, endolithic alga that lives inside marine calcium carbonate structures such as limestone reef rocks, bivalve shells, and the skeleton of corals [10]. It has a broad absorption spectrum, allowing it to use near-infrared light not absorbed by the overlying layer of Symbiodiniaceae [11]. *Ostreobium* provides nutrients and photosynthates to the coral [12], and during bleaching events, it can temporarily substitute the role of *Symbiodinium* [13]. However, with increased light availability, *Ostreobium* increases in biomass and thus in respiration, which may eventually lead to chemical dissolution of the coral skeleton, causing mortality [14]. *Ostreobium* is widespread in tropical corals worldwide, and follows a biogeographical pattern similar to that of Symbiodinium, with different clades exhibiting host-species specificity [15].

Most coral microbiome studies use 16S rRNA sequencing to profile prokaryotic diversity, but plastid-bearing microeukaryotes are also captured in these amplicons and routinely discarded by standard pipelines [16]. Retaining these reads yields critical information on plastid-bearing eukaryotes [17]. Unfortunately, Symbiodiniaceae cannot be recovered via 16S metabarcoding because dinoflagellate chloroplasts are extremely reduced and organized into small minicircles [18]. However, other plastid-bearing microeukaryotes such as apicomplexans (via the apicoplast) and phototrophic algae, including green algae, red algae, haptophytes, and diatoms, are consistently recovered in 16S rRNA datasets.

In this study, we reprocessed 16S rRNA gene datasets from 45 peer-reviewed studies spanning all major oceanic regions, coral lineages, and health states. This establishes a global framework for investigating plastid-bearing microeukaryotes across coral holobionts and provides novel insights into their biogeographic patterns and host specificity.

## Methodology

### Data acquisition and processing

A literature review identified publications exploring the anthozoan microbiome across various biotic and abiotic factors (Supplemental Table 1). Short-read amplicons were mined from NCBI’s short read archive (SRA) for all corresponding published 16S rRNA gene data until 2020. SRA accessions were downloaded using the SRA Toolkit version 2.11.0. Studies were retained in this analysis provided they: (1) were accompanied by sufficient metadata, (2) the publicly available data was submitted raw, and (3) were sequenced using Illumina MiSeq. CutAdapt [19] was used to trim primer sequences spanning different hypervariable regions by employing generalized primer definitions that accounted for ambiguous bases, allowing consistent primer removal across studies (Supplementary Table 2). Following primer trimming, amplicon sequence variants (ASVs) were inferred using DADA2 version 1.20.0 [20] and taxonomy was assigned using a combined reference database containing Silva v1.32 [21]. Sequences classified as “chloroplast” by Silva were subsequently reassigned using the PR2 database version 4.13.0 [22].

Sequencing read lengths varied across studies (2×250 bp and 2×300 bp); we retained only samples targeting the V4 or V3–V4 hypervariable regions to ensure comparable amplicon length and region coverage. Metadata was obtained from the SRA and manually curated using information from the original publications (Supplementary Table 3). The ASV table, taxonomy table and curated metadata were integrated into a phyloseq object using the R package “Phyloseq” version 1.46.0 [23]. Samples with less than 250 reads were removed, and only ASVs present within the top 99% cumulative abundance in at least one sample were retained.

### Microbiome analyses

Alpha diversity patterns were explored using the R package “microbiome” v1.24.0 [24]. Shapiro-Wilk tests indicated non-normal distributions across most categories, therefore pairwise comparisons were performed using the nonparametric Wilcoxon rank-sum test. Raw p-values were adjusted using the Benjamini-Hochberg False Discovery Rate (FDR) method. Significance was defined as *p*<0.05 after FDR correction (*), with *p*<0.01 (\*\**) and p<0.001 (*\*\*\*) considered highly significant. Community profiles, abundance plots and heatmaps were generated using the R package “ggplot2” [25]. For bubble plots, only taxa with mean relative abundance of at least 0.1% across samples were retained. UpSet plots were generated using “UpsetR” version 1.4.0 [26]. Differential abundance was assessed with “ANCOMBC” version 2.4.0 [27], with a minimum prevalence cutoff of 1%, FDR-adjusted p-values, and pairwise group comparisons. ASVs were considered significant at q < 0.05, and a log2 fold-change cutoff of 1 was applied to highlight the top differentially abundant taxa, visualized via a volcano plot.

### Phylogenetics and evolutionary placement analysis

Phylogenetic analyses were performed on the apicomplexan and *Ostreobium* sequences recovered from the dataset. To place short ASVs in a phylogenetic context, we first built maximum-likelihood reference trees for each group, then used the Evolutionary Placement Algorithm (EPA) to position ASVs onto the fixed topologies [28]. Black corals were excluded due to insufficient sampling depth.

To construct the apicomplexan 16S rRNA gene tree, a dataset of 170 apicomplexan sequences was compiled, with three rhodophyte sequences included as an outgroup. A total of 589 Ulvophyceae sequences were compiled for the *Ostreobium* reference tree, 506 of which were from the genus *Ostreobium*. For both datasets, sequences were sorted and clustered at 99.5% similarity with VSEARCH 2.30.0 [29] and aligned with MAFFT v7.525 [30]. Alignments were manually inspected in AliView v1.28 [31] and then trimmed with Trimal v1.5.0 [32], using a gap threshold of 0.3, a similarity threshold of 0.001, and a minimum conservation of 30%. Substitution models were selected with ModelTest-NG v0.1.6 [33], and maximum-likelihood trees were built with RAxML v8.2.12 [34], using 1,000 bootstrap replicates and the GTR+Γ+I model for Apicomplexa and GTR+Γ for *Ostreobium*.

Apicomplexan and *Ostreobium* ASVs from healthy coral samples were placed onto each reference topology using EPA in RAxML v8.2.12, with dataset-specific substitution models selected by ModelTest (GTR+Γ for Apicomplexa and GTR+Γ+I for *Ostreobium*). Only placements contributing at least 20% of the total likelihood weight were retained. Five *Ostreobium* ASVs (ASV_93989, ASV_100539, ASV_179338, ASV_202025, and ASV_57577) fell outside the *Ostreobium* clade and were excluded, after which EPA placement was repeated using the corrected dataset. The resultant EPA trees were visualized and annotated using the Interactive Tree of Life (iTOL) platform [35], with log-transformed heatmaps and histograms generated in R and imported into iTOL. The scripts used for data processing and analysis are available at https://github.com/delCampoLab/coral-meta.

## Results

### Shannon diversity of plastid-bearing coral symbionts

To assess broad patterns in plastid-bearing microeukaryotic diversity, we compared Shannon diversity across coral lineages, coral families, large marine ecosystems, and host health states. Among healthy corals, Shannon diversity values ranged from 1.469 ± 0.885 in robust corals to 1.858 ± 0.676 in black corals. Black corals were significantly more diverse than robust corals (Wilcoxon rank-sum test; padj = 8.3 × 10⁻³) (Supplementary Figure 1A), whereas no other pairwise comparisons were significant.

At the coral family level, Shannon diversity spanned a broader range, from 0.720 ± 0.775 in Dendrophylliidae to 2.729 ± 0.652 in Briareidae, yet no consistent pattern emerged among coral families (Supplementary Figure 1B). Similarly, Shannon diversity varied across large marine ecosystems without a clear biogeographic trend, with values ranging from 0.458 ± 0.575 in the Insular Pacific-Hawaiian region to 1.999 ± 0.741 in the Caribbean Sea (Supplementary Figure 2A).

Diseased corals showed significantly higher Shannon diversity than both healthy (Wilcoxon rank-sum test; padj = 1.25 × 10⁻⁷) and bleached corals (Wilcoxon rank-sum test; padj = 1.02 × 10⁻⁴), while no significant differences were observed between healthy and bleached (Wilcoxon rank-sum test; padj = 3.3 × 10⁻¹) (Supplementary Figure 3).

### Coral plastid bearing symbionts across regions, host lineages and health states

Coral plastid-bearing eukaryotes were dominated by diatoms, apicomplexans, rhodophytes and chlorophytes across geographic regions (Supplementary Figure 2B), coral lineages (Figure 2), and health states (Figure 2; Supplementary Figure 3B).

**Figure 1.**
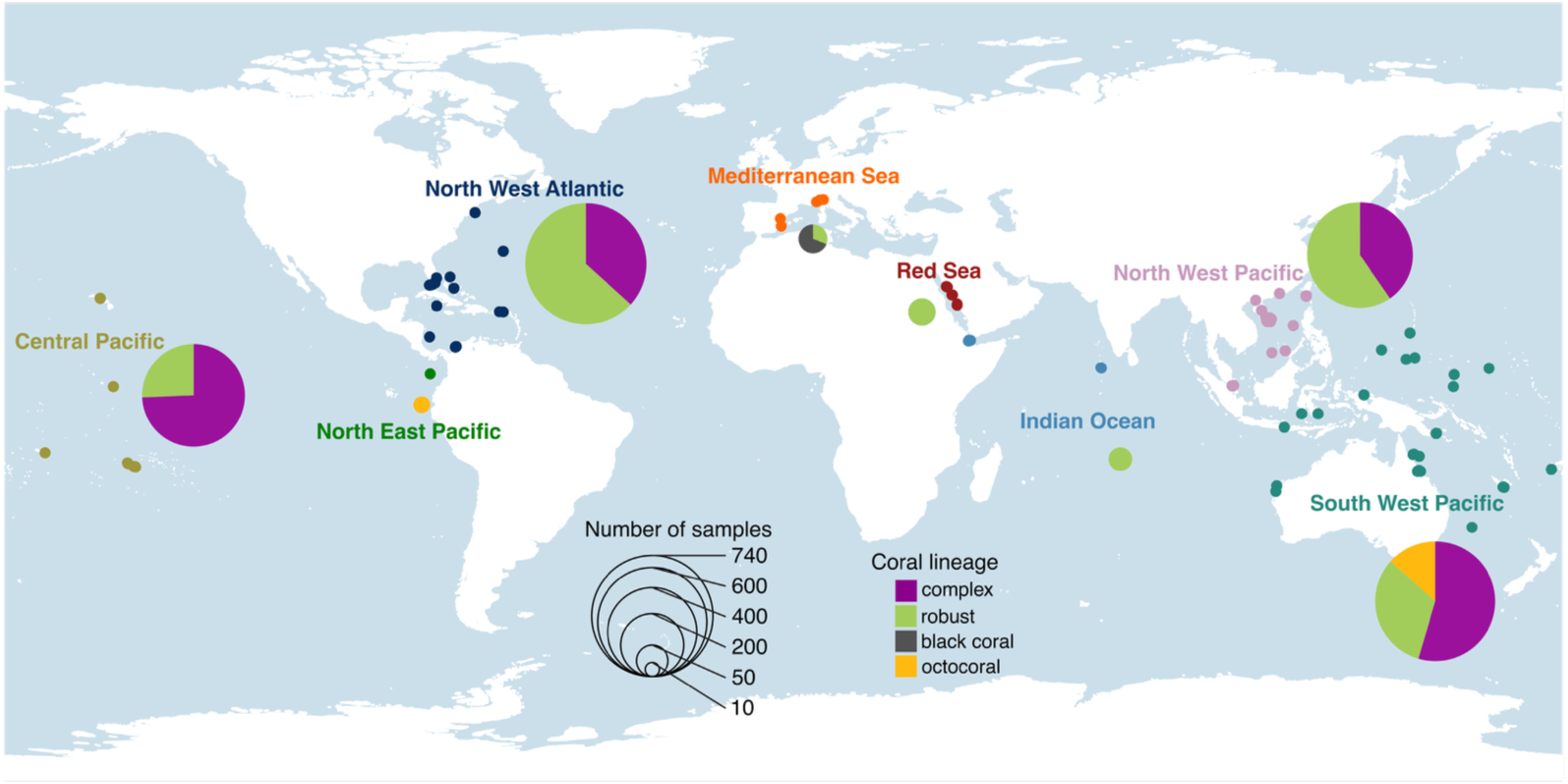
Global distribution of coral samples included in this study. Pie chart size reflects sample number; colors indicate the relative abundance of each coral taxonomic group (complex corals, robust corals, black corals, and octocorals). The map shows the same oceanic regions used by McCauley et al [55].

**Figure 2.**
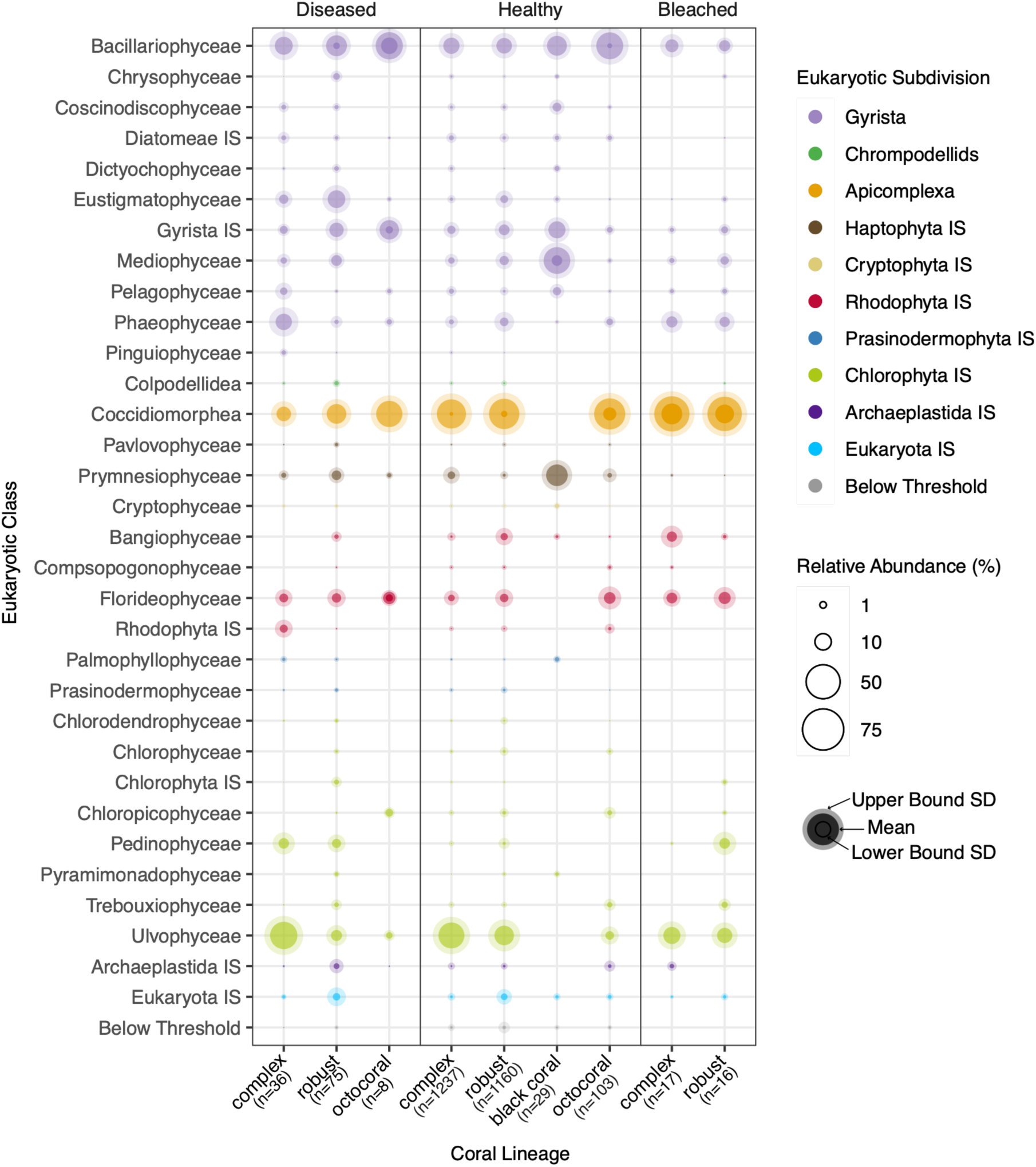
Relative abundance of plastid-bearing eukaryote classes across complex scleractinians, robust scleractinians, black corals, and octocorals, separated by health status (diseased, healthy, bleached). Bubble size represents mean relative abundance. Taxa are displayed if mean relative abundance exceeds 0.1% in at least one health state and are shown across all health states regardless of individual threshold, allowing direct cross-condition comparison. Inner and outer halos represent the lower and upper bound standard deviation (SD), respectively. Taxa below 0.1% contributions are grouped as “Below Threshold”.

Across large marine ecosystems, communities were generally dominated by Apicomplexa and Ulvophyceae (Supplementary Figure 2B). However, Apicomplexa were notably absent from several regions: the Mediterranean Sea, where communities were instead dominated by Mediophyceae, Prymnesiophyceae, and Florideophyceae; the Southeast U.S. Continental Shelf, where only Ulvophyceae were detected; and the Northeast U.S. Continental Shelf, though this region was represented by a single sample. In the West Central and Northwest Australian Shelf, Apicomplexa were detected but at markedly reduced abundances relative to other regions.

At the host lineage level, community composition was broadly similar between complex and robust corals across all health states (Figure 2). Differences between the two lineages were limited to a small number of taxa and were largely health-state dependent. For example, among diseased corals, Phaeophyceae reached higher relative abundances in complex corals than in robust corals, whereas bleached robust corals exhibited an increased relative abundance of Pedinophyceae relative to bleached complex corals.

### The structure of the coral plastid-bearing symbiont community

Despite the high overall diversity observed in coral plastid-bearing symbionts, community structure was strongly shaped by a limited number of highly prevalent ASVs across healthy (Figure 3), bleached (Supplementary Figure 4), and diseased corals (Supplementary Figure 5). In healthy corals, Corallicolida dominated the 50 most abundant ASVs (56%), followed by *Ostreobium* (22%). Other taxa included diatoms (e.g., *Thalassiosira*, *Navicula*), eustigmatophytes (*Nannochloropsis*), and occasional representatives of red and green algae. These high abundance ASVs were often host-clade specific, with distributions restricted to complex, robust, or octocoral lineages. Only two Corallicolida (ASV_534 and ASV_1774) were shared across all three coral groups. Three Corallicolida ASVs were detected exclusively in octocorals, the remaining Corallicolida were either specific to complex or robust corals, or shared between them.

**Figure 3.**
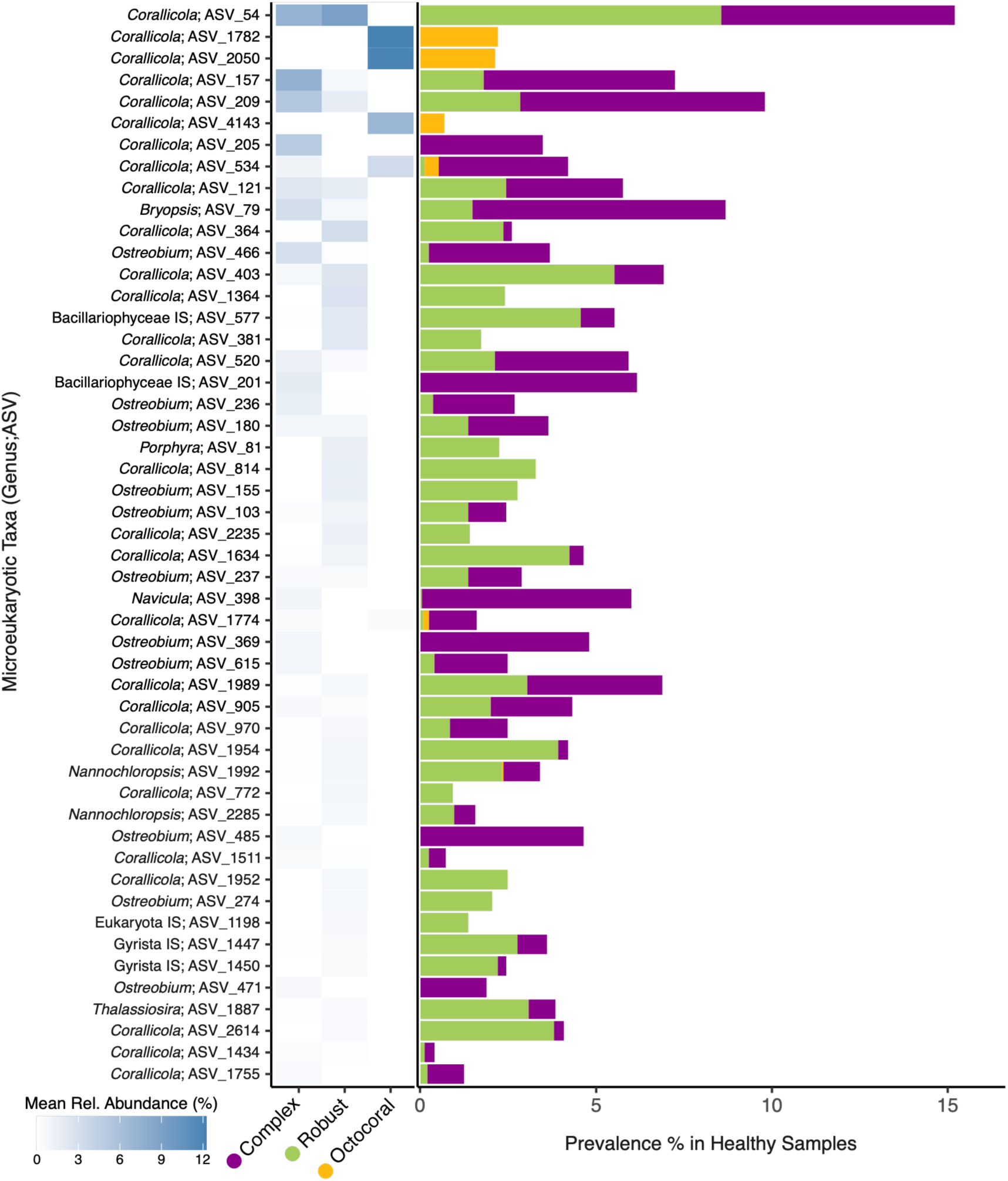
Heatmap of mean relative abundance of the most abundant ASVs in healthy corals across complex corals, robust corals, and octocorals. Color intensity reflects mean relative abundance. Sidebar bars indicate the number of healthy samples in which each ASV was detected, colored by coral lineage.

In bleached corals, the 50 most abundant ASVs remained dominated by Corallicolida but exhibited a more taxonomically diverse phototrophic fraction, including several brown algae lineages, diatoms, and chlorophytes (Supplementary Figure 4). Host specificity was observed: the most abundant ASV overall (ASV_2115, a corallicolid) was detected exclusively in complex corals, and *Ostreobium* ASVs among the dominant taxa occurred in either complex or robust corals but not both.

In diseased corals, the microbiome composition shifted more markedly. Corallicolida were reduced to 10% of the most abundant taxa, and *Ostreobium*, diatoms, and other stramenopiles increased in relative representation (Supplementary Figure 5). Despite this compositional shift, the pattern of host specificity was maintained: as in bleached corals, *Ostreobium* ASVs were restricted to either complex or robust corals.

### The specificity of the coral plastid bearing community

The pattern of host restriction extended across the dataset. Among healthy corals, most ASVs were host-specific and not shared across host categories (Figure 4A). Specifically, 6,119 ASVs were unique to complex scleractinians (85% of total complex ASVs), 2,451 ASVs to robust scleractinians (68% of total robust ASVs), 151 to black corals (88% of black coral ASVs), and 346 to octocorals (67% of octocoral ASVs) (Supplementary Figure 6). Consistent with their closer phylogenetic relatedness, the two Scleractinia lineages shared more ASVs with each other than with either black corals or octocorals, and no ASVs were shared across all four host groups.

**Fig 4.**
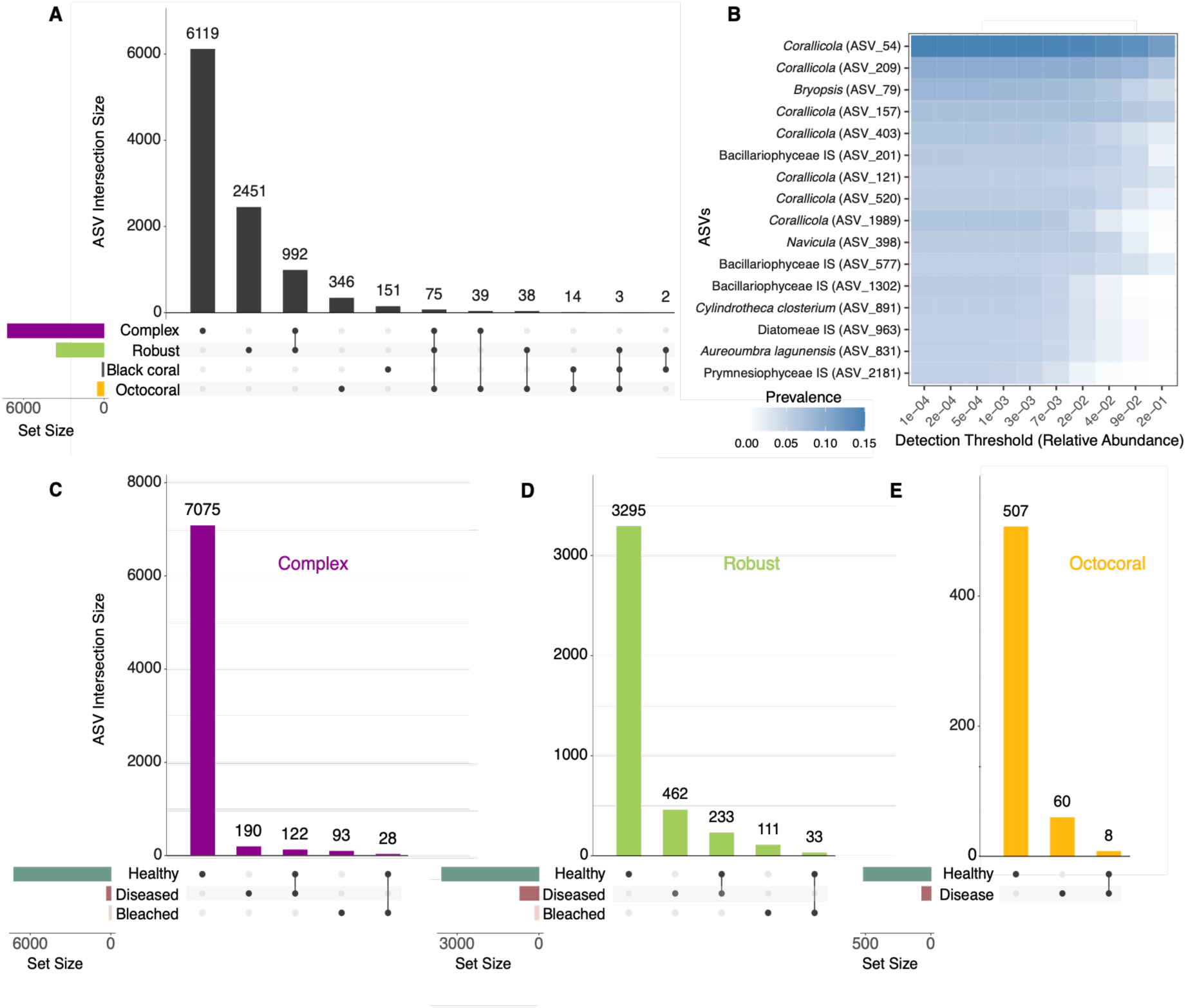
Specificity of the coral microbiome. A) UpSet plot showing the number of ASVs unique to and shared across coral hosts in healthy corals. B) Core microbiome of healthy corals (ASVs present in ≥X% of samples), colored by prevalence. C–E) ASVs unique to and shared across health states (healthy, bleached, diseased) in complex (C), robust (D), and octocorals (E).

Despite this strong host specificity, a subset of ASVs was broadly distributed across healthy corals. Using a prevalence threshold of ≥5% of samples and a minimum relative abundance of ≥0.01% per sample (detection ≥1×10⁻⁴), we identified 16 “core” ASVs (Figure 4B). These were dominated by Corallicolida (7 ASVs) and diatoms, with ASV_54, a corallicolid, being the most prevalent and abundant.

Within each host category, most ASVs were unique to a single coral health state, with only a small fraction of ASVs shared among healthy, bleached, and diseased corals (Figure 4C-E). Notably, diseased corals shared a higher percentage of ASVs with healthy corals than bleached corals did.

To identify taxa driving differences between the Scleractinia lineages, we performed differential abundance analysis using ANCOM-BC and visualized results in a volcano plot, highlighting the 10 most differentially abundant ASVs (Figure 5). ASVs enriched in complex scleractinians (negative log2 fold change) included two *Corallicola* ASVs (ASV_157, ASV_121), one *Ostreobium* ASV (ASV_615), one diatom ASV (ASV_398), and a green algae ASV (ASV_79). In contrast, ASVs enriched in robust scleractinians (positive log2 fold change) included two *Corallicola* ASVs (ASV_364, ASV_970) and three diatom ASVs (ASV_1447, ASV_577, ASV_2401).

**Figure 5.**
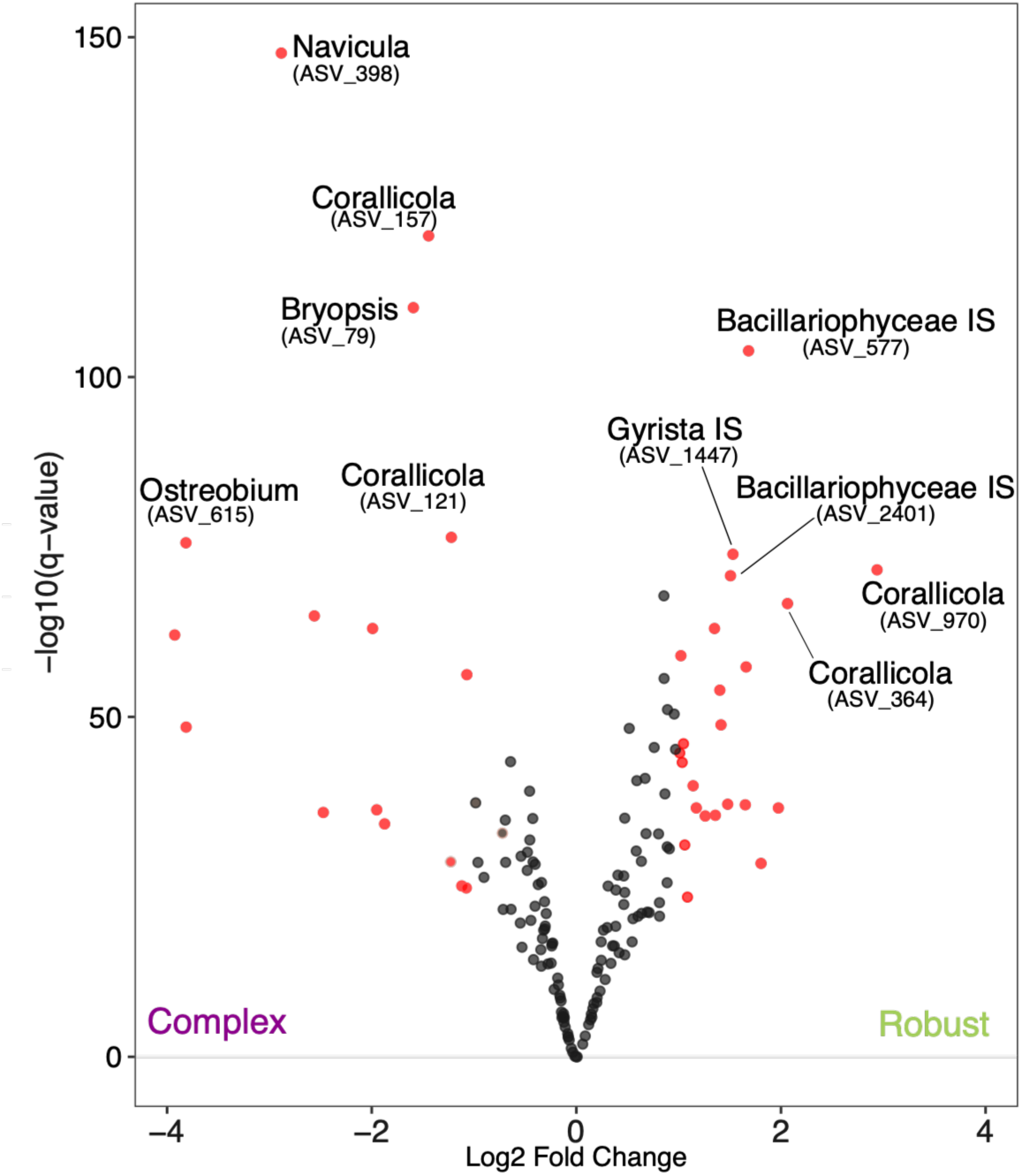
ANCOM-BC volcano plot comparing microbial ASV enrichment between complex and robust scleractinians (healthy samples only). Each point represents an ASV; red points meet both significance thresholds (q < 0.05 and |Log2 Fold Change| > 1), black points do not. The 10 ASVs with the highest significance among those meeting both thresholds are labeled with their taxonomic identifier. Negative Log2 Fold Change indicates enrichment in complex scleractinians; positive values indicate enrichment in robust scleractinians.

### Phylogenetic placement of Apicomplexa ASVs

Given the high abundance and prevalence of Apicomplexa across regions and hosts, their reads were examined further using phylogenetic placement. An apicomplexan SSU reference phylogeny was constructed from 170 apicomplexan sequences retrieved from NCBI. A total of 421 Apicomplexan ASVs from our healthy coral dataset were placed onto the apicomplexan SSU reference phylogeny using EPA (Figure 6).

**Figure 6.**
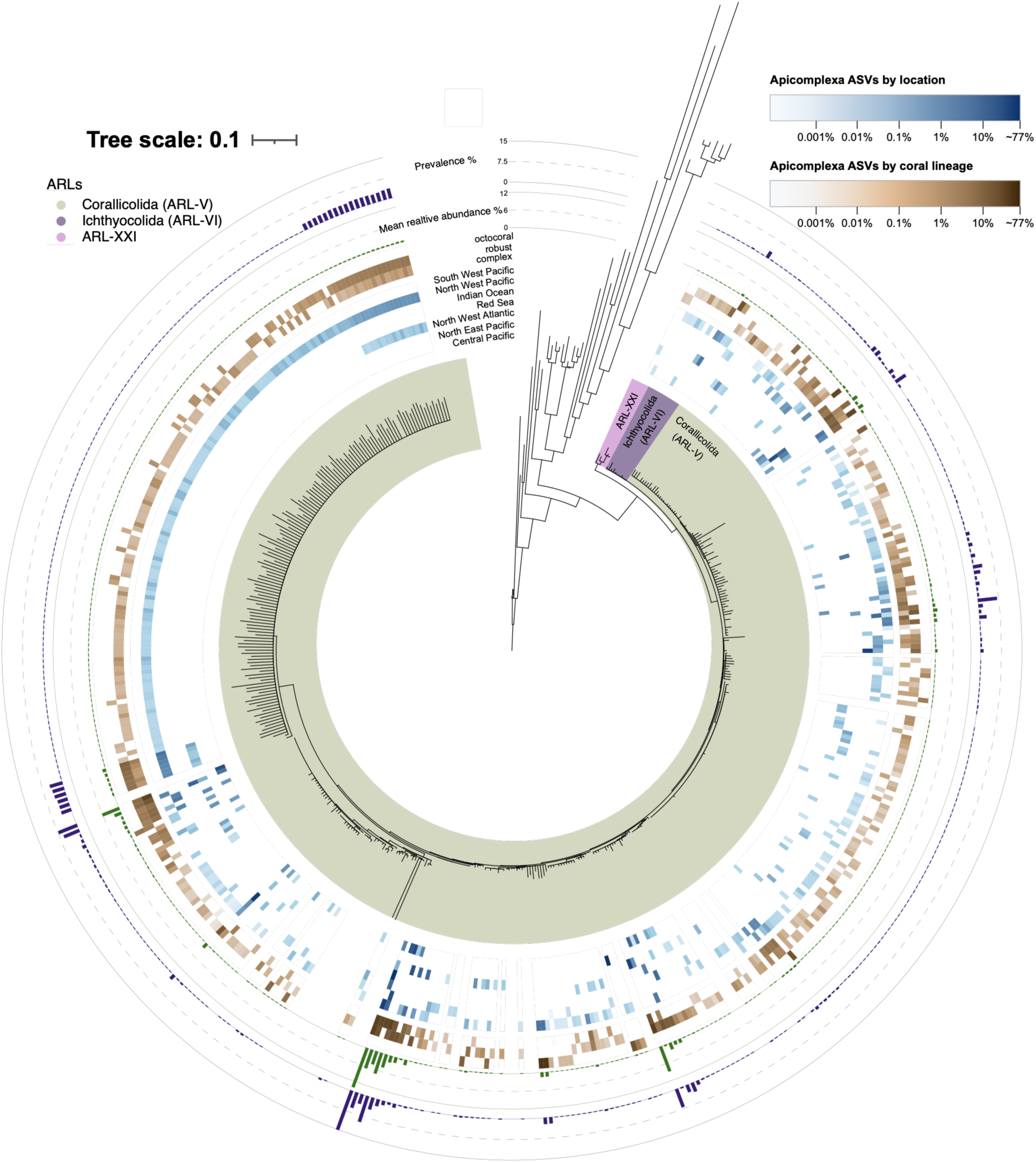
Phylogenetic classification, distribution across water masses and coral type, prevalence and relative abundance of apicomplexan reads. Maximum-likelihood reference tree of Apicomplexa, with healthy-coral apicomplexan ASVs placed via the Evolutionary Placement Algorithm. Two concentric, zero-masked heatmap rings show log10-transformed relative abundance of placed ASVs by coral host type (brown, inner ring: octocoral, robust, complex) and by oceanic region (blue, outer ring); these are followed by two bar-graph rings showing mean relative abundance (green) and prevalence (purple), both expressed as percentages, for each ASV across the dataset. Scale bar = branch length in substitutions per site.

The large majority of apicomplexan ASVs were placed within the Corallicolida clade. Exceptions included two ASVs (ASV_262986 and ASV_540285), which grouped with the Ichthyocolida [36], and a set of 10 ASVs positioned on a branch intermediate between Corallicolida and Ichthyocolida.

Coral-associated apicomplexan ASVs were detected across complex, robust, and octocoral hosts, with the highest number of ASVs observed in robust corals, followed by complex and octocorals. Phylogenetic placement revealed no clear clustering by host type, as ASVs associated with different coral host types were broadly interspersed throughout the tree. Similarly, geographic origin did not correspond to deep phylogenetic structure, with ASVs from different geographical locations distributed across multiple branches of the tree.

At finer scales, however, some regional clustering was apparent. A distinct group of closely related ASVs was exclusively detected in samples from the Northwest Pacific, forming a geographically coherent cluster within Corallicolida. Another group of related ASVs was detected in both the Northwest Pacific and Northwest Atlantic and was associated with both complex and robust corals. Despite these localized patterns, most apicomplexan ASVs showed restricted distributions, typically occurring in a single coral host type and a single geographic region. The most abundant ASV overall, ASV_54, was an exception, since it was detected in both Scleractinia lineages and across multiple regions, including the Central Pacific, the North West Atlantic, the Red Sea, the Indian Ocean, and the South West Pacific.

### Phylogenetic placement of *Ostreobium* ASVs

*Ostreobium,* was similarly selected for phylogenetic analysis. A reference SSU rRNA phylogeny was constructed using sequences from NCBI and from del Campo et al. (2017). We identified 3,422 *Ostreobium* ASVs in our healthy corals dataset and placed them on the SSU reference phylogeny using EPA (Figure 7). The resulting phylogeny recovered the three major *Ostreobium* clades (OST1, OST2 and OST3) previously described by del Campo et al. (2017) and additionally revealed three smaller clades, here designated OST4, OST5, and OST6 (Figure 7).

**Figure 7.**
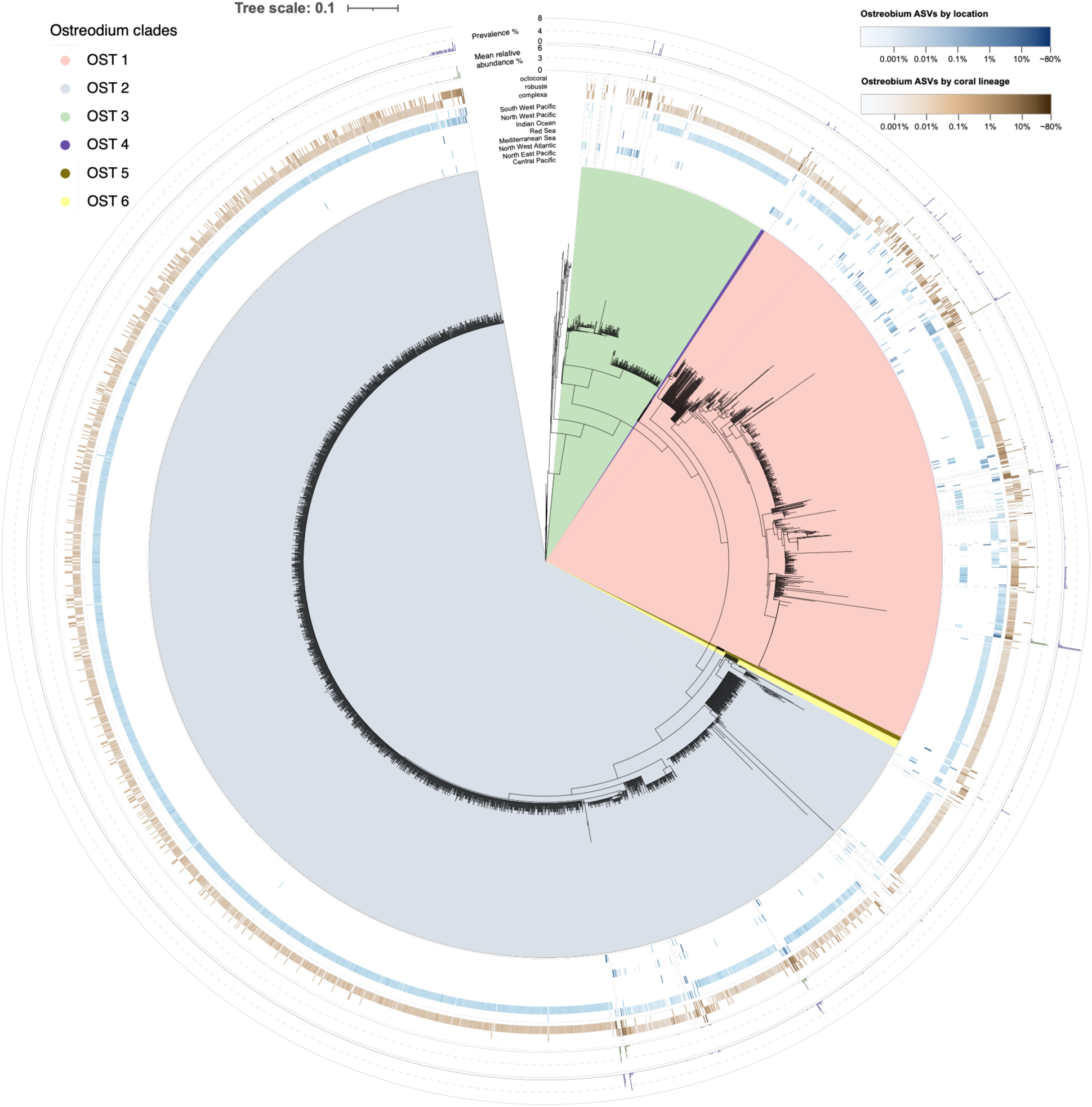
Phylogenetic classification, distribution across water masses and coral type, prevalence and relative abundance of *Ostreobium* reads. Maximum-likelihood reference tree of Ulvophyceae, with healthy-coral *Ostreobium* ASVs placed via the Evolutionary Placement Algorithm. Colored sectors denote the six recovered clades: OST1 (yellow), OST2 (green), OST3 (purple), and the smaller OST4–OST6 clades. Two concentric, zero-masked heatmap rings show log10-transformed relative abundance of placed ASVs by coral host type (brown, inner ring) and by oceanic region (blue, outer ring), followed by bar-graph rings showing mean relative abundance (green) and prevalence (purple), both as percentages. Scale bar = 0.1 substitutions per site.

Overall, *Ostreobium* ASVs were most commonly associated with complex corals, followed by robust corals, and least commonly with octocorals, both in absolute numbers and mean relative abundance. Geographically, the North West Pacific contained the greatest number of ASVs (3,409), followed by the North West Atlantic (1,472), the South West Pacific (939), and the Central Pacific (369). ASVs were also detected in the Indian Ocean, the North East Pacific, and the Red Sea, though in considerably smaller numbers. Notably, the North West Pacific ranked only third in sample size (n=539) yet yielded the highest *Ostreobium* ASV richness. In contrast, the South West Pacific, which had the greatest number of samples, contained substantially fewer ASVs (939).

Phylogenetic structure further reflected coral host and regional partitioning: OST2, which contained the highest number of ASVs, and was dominated by sequences associated with complex corals from the Northwest Pacific, with a smaller but distinct cluster of ASVs associated with robust corals from the Southwest Pacific. More broadly, most ASVs across the phylogeny were restricted to a single host type and were typically detected in only one geographic region, mirroring the host and geographic specificity described for Corallicolida above.

## Discussion

This study offers the most comprehensive view to date of plastid-bearing microeukaryotic symbiont diversity and distribution across a broad range of coral taxa, geographical locations, and health states.

### Plastid-bearing microeukaryotes in corals show high diversity but alpha diversity is not explained by host lineage, geography, or health state

Healthy corals harbored a diverse plastid-bearing symbiont community (53 classes in total, 32 consistently above 0.1% mean relative abundance) yet Shannon diversity showed weak structuring across host phylogeny, geography, or health state. Black corals showed significantly higher diversity than the two Scleractinia lineages; however, this pattern did not hold at finer taxonomic resolution. Shannon diversity varied widely within coral families and across large marine ecosystems, with no consistent trend in either case. In contrast to bacterial microbiomes, where richness declines with latitude [37], no such pattern emerged here. The absence of strong diversity gradients suggests that plastid-bearing symbiont diversity is shaped by multiple interacting factors rather than a single driver such as host phylogeny or geography.

In terms of health, diseased corals exhibited significantly higher Shannon diversity than both healthy and bleached corals, a pattern consistent with bacterial coral microbiome studies [38, 39]. Increased diversity in diseased corals may reflect a breakdown of host-mediated filtering mechanisms, allowing colonization by opportunistic biofilm colonizers such as diatoms, which are normally less abundant in healthy tissues.

The coral plastid-bearing symbiont community was dominated by diatoms, apicomplexans, rhodophytes and chlorophytes, pattern also observed by del Campo et al on their recent global survey of the coral-associated eukaryome [40]. ASVs of Corallicolida and *Ostreobium* were consistently detected across hosts and locations, confirming that they are the most widespread and abundant coral plastid-bearing symbionts after the Symbiodiniaceae [7, 15]. Lower relative abundances of Corallicolida were observed in samples from the Northwest and Central West Australian shelf compared to other regions; however, these locations were represented exclusively by 13 samples of *Stylophora pistillata* collected at Ningaloo Reef. This likely reflects the eukaryome composition of this particular species rather than a true regional absence of Corallicolida. Similarly, Mediterranean samples comprised only *Cladocora caespitosa* and *Antipathella subpinnata*, neither of which has been previously reported to harbor Corallicolida, explaining the absence of Corallicolida from our Mediterranean samples.

This dominance of Corallicolida across the dataset was also reflected at the ASV level: among the 50 most abundant ASVs in healthy corals, most were Corallicolida, followed by *Ostreobium*. However, only a handful of ASVs consistently reached high prevalence across samples, indicating that the broader Corallicolida and *Ostreobium* diversity is unevenly distributed, with most of the signal concentrated in a small number of dominant ASVs. Other recurring taxa included diatoms (e.g., *Nannochloropsis*, *Thalassiosira*, *Navicula*), which may represent epibionts or ingested prey rather than stable symbionts [41, 42]. The most prevalent genera of diatoms in the open ocean include *Chaetoceros*, *Thalassiosira*, *Actinocyclus* and *Pseudo-nitzschia* [43]. Of these, only *Thalassiosira* and *Chaetoceros* appeared in our dataset, suggesting coral-specific associations. Notably, *Thalassiosira* appeared consistently across most sampling locations and ranked among the most abundant ASVs, pointing to either a recurrent association with coral tissue or a dietary preference for this particular diatom genus. Red and green algae were also present across hosts and locations, with Florideophyceae being the most abundant and widespread red algal class, consistent with recent work on plastid symbionts in the Southwestern Atlantic [44].

The dominant community in bleached corals was broadly similar to that in healthy corals, consistent with bleaching representing a disruption of the coral-Symbiodiniaceae symbiosis rather than a wholesale restructuring of the broader microbiome. Despite Ostreobium being known to proliferate in bleached corals due to increased light availability [13], we did not observe a clear increase in its relative abundance. This could reflect temporal sampling bias (early-stage bleaching before proliferation begins), or clade-specific responses, as *Ostreobium* clades differ in their spectral niche preferences and photoacclimation responses [45], with some lineages potentially proliferating while others decline, leaving aggregate abundance unchanged.

Coral disease represents a more severe disturbance than bleaching, often involving infection, tissue loss, and host cellular damage, among other impacts [46]. Diseased corals showed a more marked compositional shift than bleached ones. The most notable change was a reduction in Corallicolida among the most abundant ASVs: given that Corallicolida are intracellular symbionts, we hypothesize that the tissue degradation and host cell lysis associated with disease may diminish their abundance, though this awaits experimental validation. This decline in Corallicolida was accompanied by an increase in diatoms, consistent with their role as opportunistic colonizers of compromised tissue [42].

### Coral plastid-bearing symbionts are highly host-specific, with a conserved core dominated by Corallicolida

Host specificity was clearly evident across analyses: upset plots showed minimal overlap between host groups, and heatmaps revealed that the most abundant ASVs were in most cases restricted to either scleractinian corals or octocorals.

Not all cases can be interpreted as true host preference, as some trends may reflect variations in sampling effort. The *Ostreobium* ASV_155, for instance, was found only in robust corals of the Northeast US Continental Shelf, but only robust corals were sampled in that region, leaving host specificity untested. In other cases, however, multiple coral lineages co-occur at the same site and microbial ASVs show preference for a specific lineage, indicating potential signs of phylosymbiosis, where host evolutionary shapes the associated microbiome. The clearest example is the Corallicolida ASV_54, the most abundant and widespread ASV in the dataset, which was recovered from major marine ecosystems across the globe (the Caribbean Sea, the Red Sea, the Arabian Sea, the Northeast, Central, and Northwest Australian Shelf, the Indonesian Sea, Micronesia, and Polynesia) yet was entirely absent from octocorals. Although ASV_54 was classified as part of the core microbiome, since it was undetected in octocorals, its core status reflects the scleractinian-dominated composition of the dataset rather than a true pan-coral distribution.

By contrast, three abundant corallicolids (ASV_1782, ASV_2050, ASV_4143), all recovered from the Northeast Australian Shelf where complex, robust, and octocorals co-occur, were found exclusively in octocorals, again showing high coral lineage specificity. Among the most abundant ASVs, the only two shared across all three host lineages, ASV_534 and ASV_1774, were widespread across locations, suggesting a more generalist ecology.

Host specificity, though more obvious in the split between scleractinian corals and octocorals, also held at a finer scale between complex and robust corals. Although community composition was broadly similar at higher taxonomic levels, many ASVs were not shared between them, and this divergence was detectable even where hosts co-occurred. The corallicolid ASV_1364, for instance, was restricted to robust corals despite being recovered from Polynesia and the Northeast Australian Shelf, sites where all three coral groups co-occurred. The same logic applied to ASV_201 (Bacillariophyceae), restricted to complex corals at the Northeast Australian Shelf under identical sampling conditions. Geography cannot account for either pattern; host identity is the common factor.

Volcano plots further supported this within-scleractinian partitioning, revealing ASVs differentially enriched in complex versus robust corals. Notably, both lineages harbored clade-enriched corallicolid ASVs, indicating that despite the overall abundance and widespread distribution of Corallicolida, their fine-scale diversity is structured by host phylogeny, a signal visible only at the ASV-resolution. The diatom Navicula ASV_398, enriched in complex corals, may reflect dietary or assimilation differences between the two lineages.

### Phylogenetic placement reveals broad host and geographic distribution of Corallicolida with localized regional structure

Phylogenetic placement confirmed that the vast majority of apicomplexan ASVs belong to Corallicolida, with only a few exceptions. Two ASVs grouped with the Ichthyocolida, a group of fish-infecting apicomplexans [36]. This most likely reflects incidental detection, either through fish feces deposited on the coral surface, or through gnathiid isopods harboring in coral colonies. Like fireworms, which act as a reservoir and potential vector dispersing Corallicolida among corals via feces [47], gnathiid isopods and leeches host Ichthyocolida and may similarly serve as vectors [48, 49].

More notably, 10 ASVs occupied a branch intermediate between Corallicolida (ARL-V) and Ichthyocolida (ARL-VI), a phylogenetic position corresponding to no currently described Apicomplexan Related Lineage (ARL) [50]. These 10 ASVs may therefore represent a novel, undescribed ARL (tentatively ARL-XXI, following the numerical system extended to ARL-XX by Janouškovec et al [51]), a finding that warrants further investigation using longer marker sequences and broader taxonomic sampling.

The corallicolid part of the tree showed no strong phylogenetic structuring overall, though host- and location-associated clustering was apparent in specific parts of the tree. One large cluster of ASVs, low in overall abundance and prevalence, was exclusive to the NW Pacific and largely restricted to complex corals. Sampling bias does not account for this: complex corals were actually underrepresented in NW Pacific samples relative to robust ones (228 vs. 336 samples), indicating a genuine host association rather than an artifact of uneven sampling. This pattern may reflect local diversification of a Corallicolida clade within the complex coral lineage. In contrast, the most abundant Corallicolida ASVs were broadly distributed across locations and host types, suggesting a distinction between a widely distributed core of dominant lineages and a periphery of geographically or host-restricted, low-abundance ASVs.

### Phylogenetic placement of *Ostreobium* reveals host-specific and geographically structured clades

*Ostreobium* displayed clearer phylogenetic structure than Corallicolida, with a net preference for complex over robust corals across the genus that was driven unevenly by individual clades. For example, OST2, the most ASV-rich clade, showed the strongest expression of this trend, with ASVs almost exclusively restricted to complex corals of the North West Pacific. The fact that the North West Pacific had a lower number of samples than the South West Pacific but a much higher number of *Ostreobium* ASVs exposes a non-linear relationship between sampling effort and ASV diversity, and suggests that the North West Pacific may be a richer reservoir of *Ostreobium* diversity.

OST1 and OST3 showed similar general trends to OST2 but with more frequent departures, including notable representation in the North West Atlantic and, in OST3, a more abundant subcluster shared between complex and robust corals. This indicates that clade identity in *Ostreobium* may partly predict host or geographic association, confirming the observations from del Campo et al 2017. One possible explanation for the overall complex/robust preference is that it reflects underlying biochemical differences between the two Scleractinia lineages: robust corals retain a complete histidine biosynthetic pathway, which is absent in complex corals [52]. While limited to a single pathway and untested for *Ostreobium* specifically, this biochemical divergence illustrates that the two lineages differ in ways plausibly relevant to symbiont compatibility, offering a testable hypothesis for the host preference observed here.

This pattern, in which host structuring emerges unevenly across individual clades rather than uniformly across a whole symbiont lineage, is not unique to *Ostreobium*. This fine-scale host specificity between complex and robust corals has previously been observed in the prevalent bacteria *Endozoicomonas* [37]. *Endozoicomonas* subdivides into host-specific subclades that associate with either complex or robust lineages, with host-swapping being a rare event throughout the evolution of scleractinian corals. This pattern is mirrored here at the ASV level across multiple symbiont taxa. Symbiodiniaceae, by contrast, show biogeographical structuring and host specificity [53], but not in a way that maps onto the complex/robust divide. Recent work testing this directly across 39 scleractinian species found that corals exhibit host specificity but lack phylosymbiosis, with *Cladocopium* and *Durusdinium* showing different cophylogenetic patterns relative to their hosts [54]. These contrasts suggest that the complex/robust divide shapes different symbiont groups differently, and at different taxonomic scales, depending on each symbiont’s biology; there is no single universal axis of coral-microbe specificity.

As with any large-scale meta-analysis, we must interpret our findings in light of several methodological constraints. This work integrates 45 source studies that differ in DNA extraction protocols, primer choice within the V3–V4/V4 regions, sequencing depth, and sample preservation, introducing technical variation that is in some cases difficult to separate from true biological signal. Sampling effort was also uneven across geography, host taxa, and health states: some regions, non-scleractinian hosts, and the bleached and diseased datasets are represented by relatively few samples. This limits statistical power and means that apparent absences cannot be confidently distinguished from true ecological absence, under-sampling, or low sequencing depth. With these caveats in mind, the breadth of the dataset nonetheless allowed us to identify robust patterns in coral-associated plastid-bearing symbiont communities.

By recovering reads typically discarded from 16S rRNA gene surveys, we showed that standard pipelines routinely overlook meaningful eukaryotic signal. Our findings show that host identity and geography strongly structure plastid-bearing symbiont communities, with Corallicolida and Ostreobium as the dominant players after the Symbiodiniaceae, confirming previous observations. Key novel findings include three previously undescribed *Ostreobium* clades, a potential diversity hotspot in the North West Pacific, and a set of apicomplexan ASVs tentatively designated ARL-XXI that warrant follow-up with longer markers and broader taxonomic sampling. Whether factors driving fine-scale host preferences reflect biochemical compatibility, co-evolutionary history, ecological filtering, or a combination remains an open question to be resolved through targeted experimental and metagenomic approaches. Taken together, these results show that plastid-derived reads are not noise to be filtered out, but a valuable window into the hidden diversity of the coral holobiont.

## Author contributions

JdC conceptualized the project. BAW conducted the literature review. BAW and JK performed the analyses. JK prepared the figures and wrote the first draft of the manuscript. JdC and BAW reviewed and edited the manuscript. JdC supervised the study and acquired funding.

## Acknowledgements

We are grateful to the authors of all publications whose data contributed to this study, and to Anthony Bonacolta for his feedback and support throughout the analysis. The AI-assisted tool Claude was used to support code review during data analysis and to correct grammar during writing.

## Data availability

All datasets analysed during the current study are publicly available in the NCBI Sequence Read Archive; their accession numbers and original sources are listed in Supplementary Table 1.

