## Supplementary Figure 1 for "Global distribution and host specificity of plastid-bearing microeukaryotes across coral holobionts"

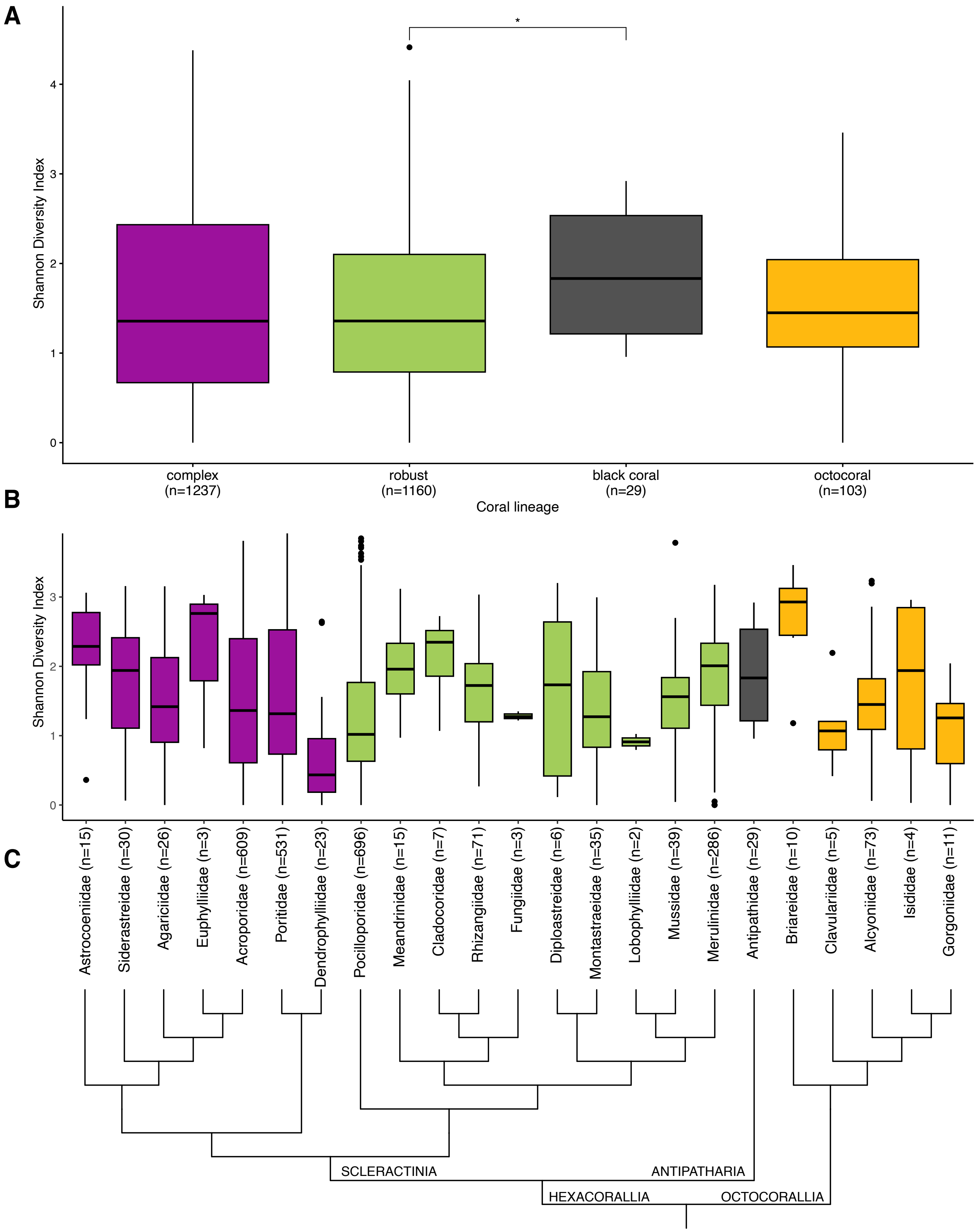


**Supplementary Figure 1. Alpha diversity across coral taxonomic groups**. A) Shannon diversity index of complex and robust corals, black corals and octocorals. Asterisks denote significance after pairwise Wilcoxon rank-sum tests with Benjamini-Hochberg FDR correction: * $p < 0.05$, ** $p < 0.01$, *** $p < 0.001$. B) Shannon diversity index of coral families. Colors correspond to figure A categories. Significance brackets are omitted from panel B due to the large number of pairwise comparisons. C) Cladogram of the coral families included in the plot above.
