## Supplementary Figure 2 for "Global distribution and host specificity of plastid-bearing microeukaryotes across coral holobionts"

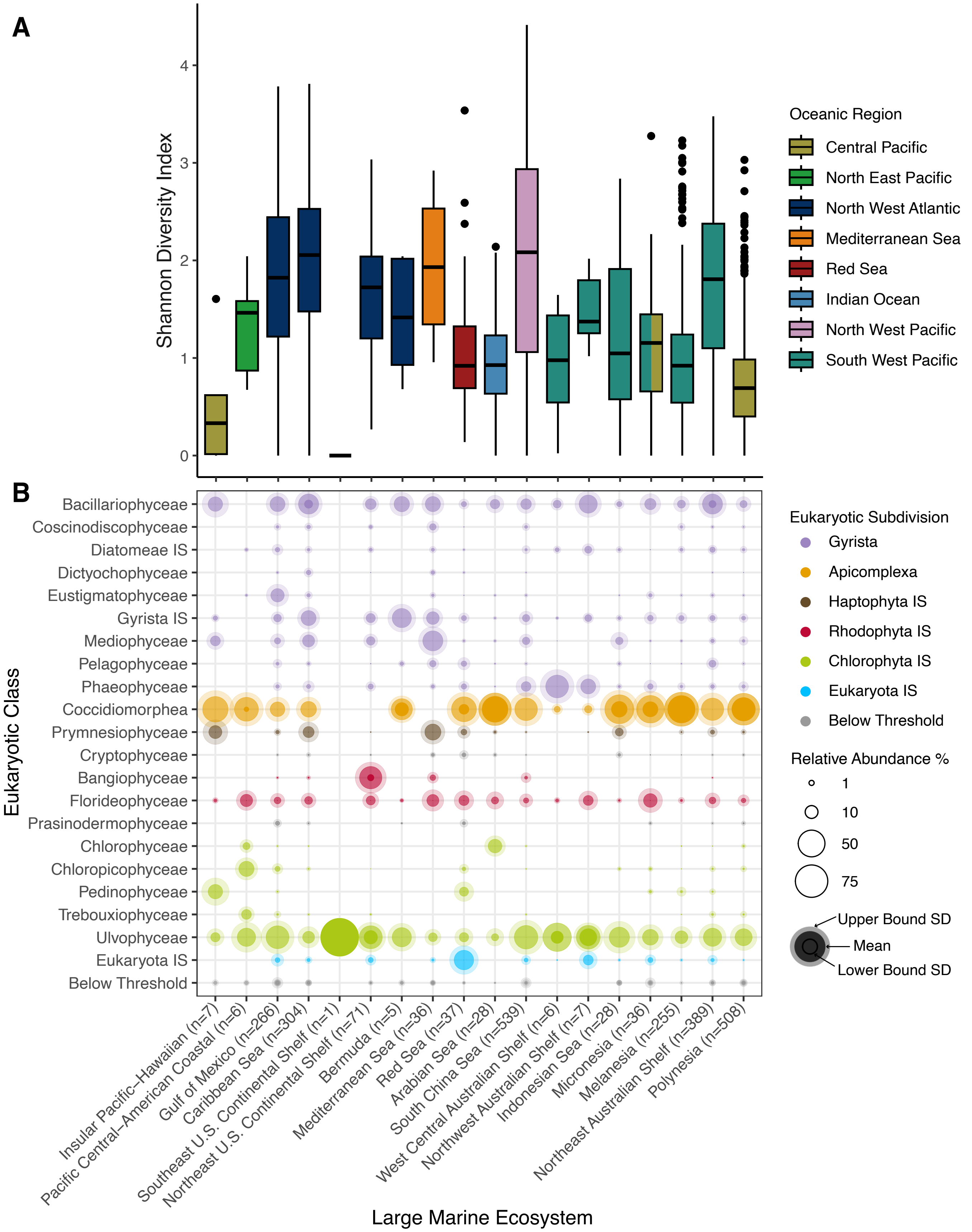


**Supplementary Figure 2**. A) Shannon diversity index of plastid bearing symbionts across large marine ecosystems, including only healthy samples. Colors of the bars correspond to different oceanic regions. B) Microbiome composition of healthy samples across large marine ecosystems, showing taxa above a mean relative abundance of 0.1%. Bubble size represents mean relative abundance, and inner and outer halos represent the lower and upper SD, respectively. Taxa below 0.1% contributions are grouped as "Below Threshold".
