## Supplementary Figure 3 for "Global distribution and host specificity of plastid-bearing microeukaryotes across coral holobionts"

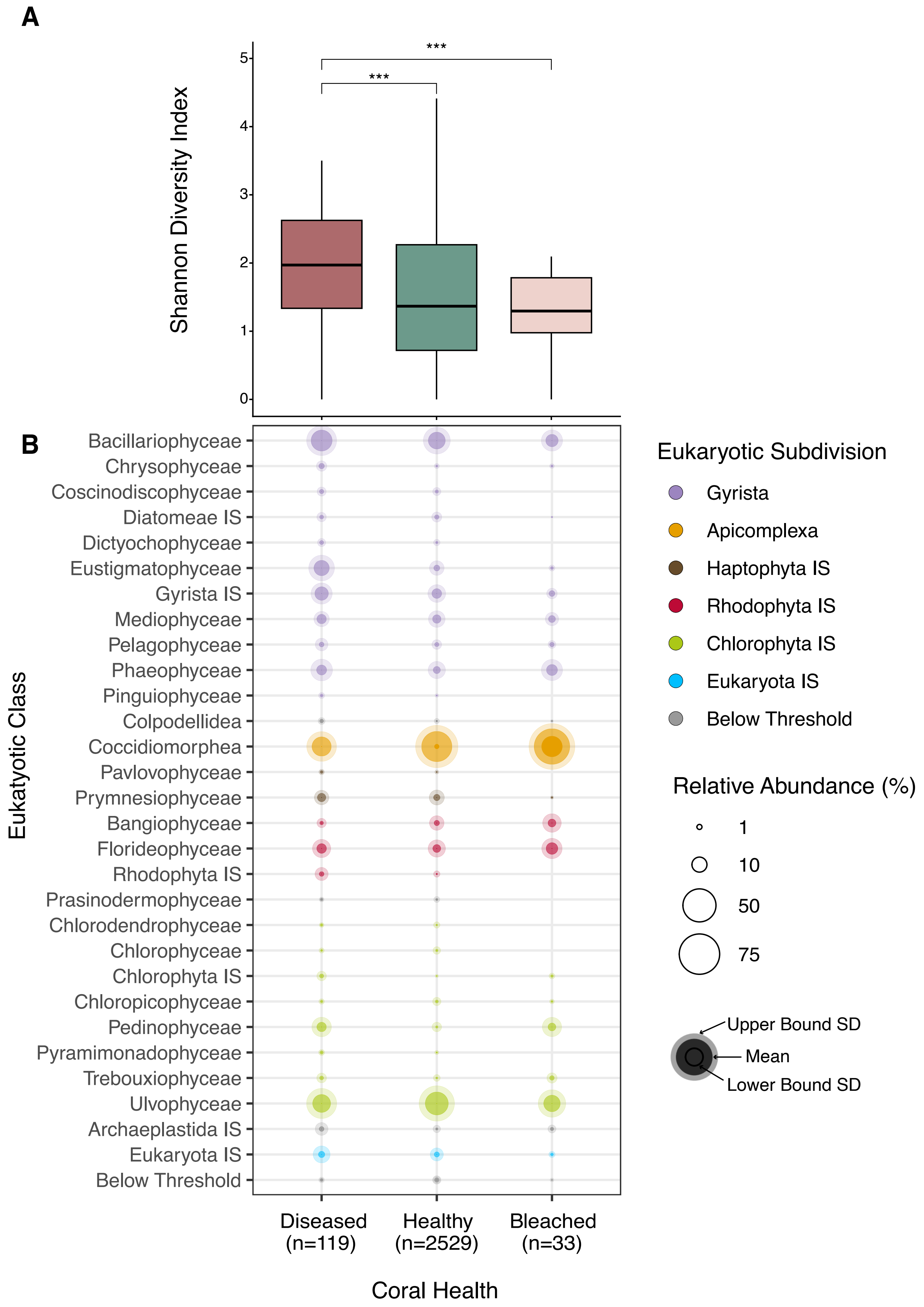


**Supplementary Figure 3**. A) Shannon diversity index of plastid bearing symbionts across coral health states. Asterisks denote significance after pairwise Wilcoxon rank-sum tests with Benjamini-Hochberg FDR correction: * $p < 0.05$, ** $p < 0.01$, *** $p < 0.001$. B) Microbiome composition across coral health states, showing taxa above a mean relative abundance of 0.1%. Bubble size represents mean relative abundance, and inner and outer halos represent the lower and upper SD, respectively. Taxa below 0.1% contributions are grouped as “Below Threshold”.
