## Supplementary Figure 4 for "Global distribution and host specificity of plastid-bearing microeukaryotes across coral holobionts"

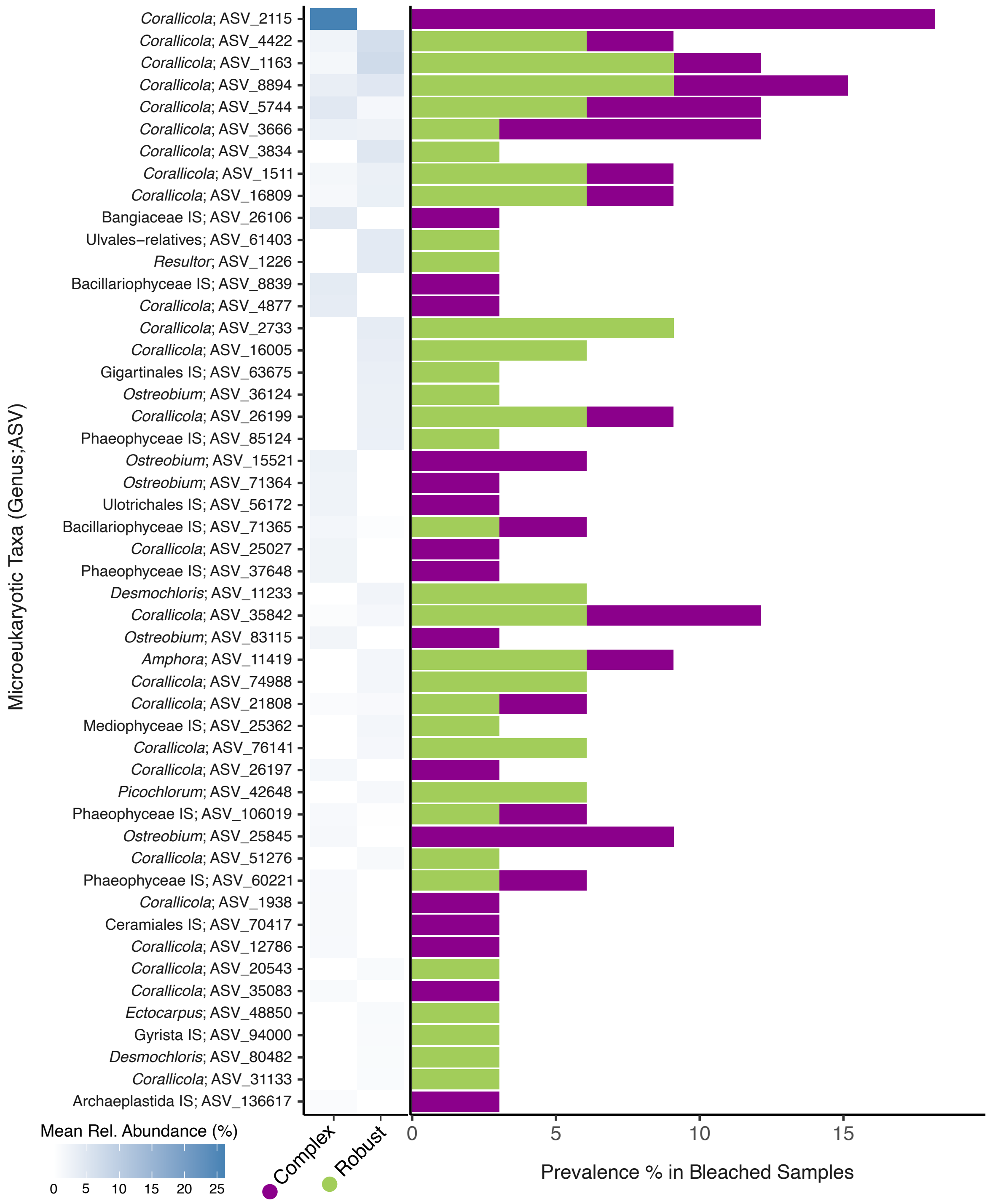


**Supplementary Figure 4.** Heatmap of mean relative abundance of the most abundant ASVs in bleached corals across complex corals, robust corals, and octocorals. Color intensity reflects mean relative abundance. Sidebar bars indicate the number of healthy samples in which each ASV was detected, colored by coral lineage.
