## Supplementary Figure 6 for "Global distribution and host specificity of plastid-bearing microeukaryotes across coral holobionts"

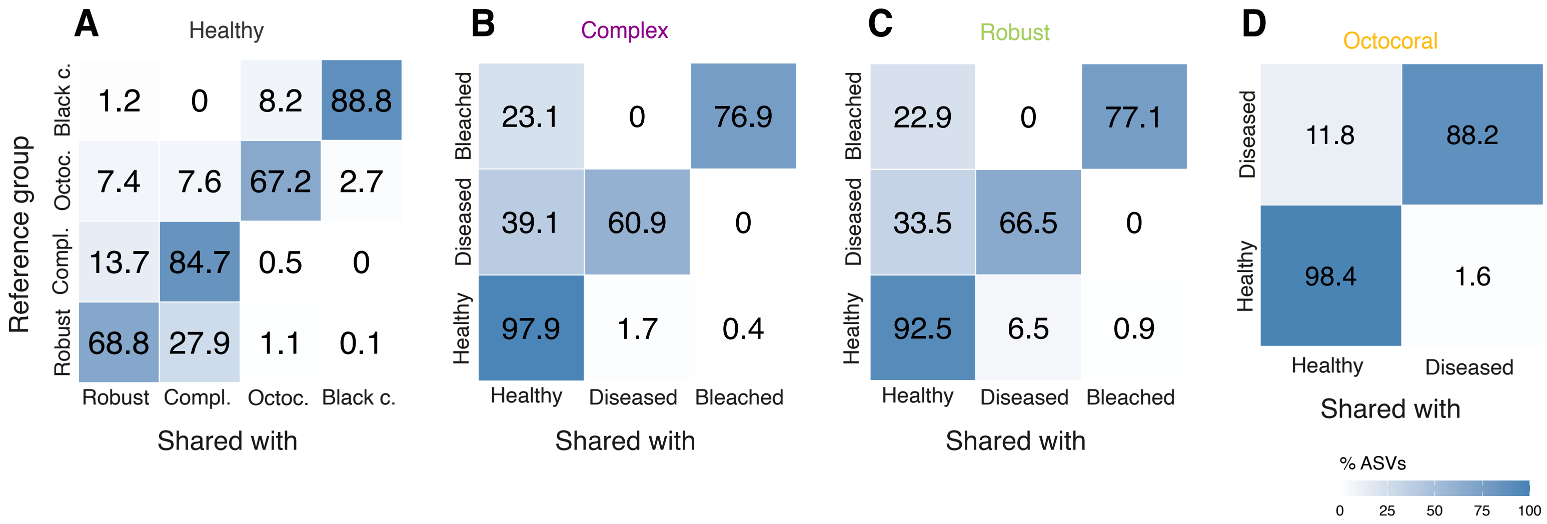


**Supplementary Figure 6. Percentage of ASVs shared between groups within each coral category, complementary to the upset plots in Figure 4.** Each cell shows the percentage of ASVs belonging to the reference group (y-axis) that are also present in the comparison group (x-axis). A) Healthy corals: ASV overlap across coral lineages (robust, complex, octocoral, and black coral). B) Complex corals: ASV overlap across health states (healthy, disease, and bleached). C) Robust corals: ASV overlap across health states (healthy, disease, and bleached). D) Octocorals: ASV overlap across health states (healthy and disease). Color scale indicates the percentage of shared ASVs, ranging from 0% (white) to 100% (blue).
